# *Schistosoma mansoni* lysine specific demethylase 1 (SmLSD1) is a druggable target involved in parasite survival, oviposition and stem cell proliferation

**DOI:** 10.1101/2020.09.17.301184

**Authors:** G. Padalino, C. A. Celatka, H. Y. Rienhoff, J. H. Kalin, P. A. Cole, D. Lassalle, C. Grunau, I. W. Chalmers, A. Brancale, K. F. Hoffmann

## Abstract

Schistosomiasis is a chronically-debilitating neglected tropical disease (NTD) that predominantly affects people living in resource-poor communities of tropical and subtropical countries. *Schistosoma mansoni*, one of three species responsible for most human infections, undergoes strict developmental regulation of gene expression that is carefully controlled by both genetic- and epigenetic- processes. As inhibition of *S. mansoni* epigenetic machinery components has been shown to impair key transitions throughout the parasite’s digenetic lifecycle, this knowledge is currently fuelling the search for new epi-drug - based anthelmintics.

In this study, the anti-schistosomal activity of 39 re-purposed *Homo sapiens* Lysine Specific Demethylase 1 (HsLSD1) inhibitors was investigated on key life cycle stages associated with both definitive (schistosomula, juvenile worms, sexually-mature adults) and intermediate host (miracidia) infection. The most active compound (compound **33**; e.g. schistosomula phenotype EC_50_ = 4.370 µM; adult worm motility EC_50_ = 2.137 µM) was subsequently used to provide further insight into the critical role of *S. mansoni* lysine specific demethylase 1 (SmLSD1) in adult worm oviposition and stem cell proliferation. Here, compound **33** treatment of adult schistosomes led to significant defects in egg production, intra-egg vitellocyte/ovum packaging and gonadal/neoblast stem cell proliferation. A greater abundance of H3K4me2 marks accompanied these phenotypes and supported specific inhibition of SmLSD1 in adult schistosomes by compound **33**. *In silico* screening indicated that compound **33** likely inhibits SmLSD1 activity by covalently reacting with the FAD cofactor.

This work suggests that evaluation of HsLSD1 - targeting epi-drugs could have utility in the search for next-generation anti-schistosomals. The ability of compound **33** to inhibit parasite survival, oviposition, H3K4me2 demethylation and stem cell proliferation warrants further investigations of this compound and its epigenetic target. This data further highlights the importance of histone methylation in *S. mansoni* lifecycle transitions.

**Author summary:** Affecting over 200 million people, schistosomiasis is a chronic disease caused by the parasitic worm *Schistosoma mansoni*. The frontline drug for schistosomiasis treatment is praziquantel. Owing to the concern surrounding praziquantel insensitivity or resistance developing, current research is directed towards the identification of novel drugs. We have focused our search for compounds that affect essential aspects of schistosome biology including parasite movement, fertility, cell proliferation and survival. Since all of these functions are potentially influenced by epigenetic regulation of gene expression, we investigated the activity of compounds that alter histone methylation status. In this report, we show that *S. mansoni* Lysine Specific Demethylase 1 (SmLSD1), a histone demethylase, is critical to miracidia-to-sporocyst transitioning, adult worm motility, egg production and parasite survival. Inhibition of SmLSD1 with compounds developed to inhibit the human paralog show promising potential as novel anti-schistosomal agents.

## Introduction

Praziquantel (PZQ) is the only drug approved for the treatment of schistosomiasis, a Neglected Tropical Disease (NTD) caused by infection with *Schistosoma* blood fluke parasites [1, 2]. Due to obvious limitations of a mono-chemotherapeutic control strategy [3-6], novel compounds with distinct mechanisms of action have been sought by researchers within both academic [7-9] as well as industrial laboratories [10] for the development of alternative or combinatorial anti-schistosomals.

In this regard, two anti-schistosomal drug discovery strategies stand out. One involves the ‘re-purposing’ of approved drugs for new indications [11, 12]. A second strategy involves the *de novo* design of drugs using either a ligand- or target-based molecular modelling approach [13, 14]. Considering that epigenetic pathways play an important role in regulating schistosome phenotype [15], controlling development [16] and responding to environmental stresses [17, 18], pharmacologic inhibition of the key protein components of these epigenetic regulators by re-purposed or *de novo* designed compounds clearly defines a promising control strategy [11, 15, 19, 20].

Using both combinatorial chemistry and drug re-purposing, we and others have pursued the investigation of protein methylation components as next generation drug targets for schistosomiasis [19-22] due to growing evidence of their impact on schistosome development and reproduction [23]. In the current investigation, we further explore the *S. mansoni* Lysine Specific Demethylase 1 (SmLSD1, Smp_150560) [19, 21] as a drug target using both late- and early-stage chemical entities developed to inhibit *Homo sapiens* LSD1 (HsLSD1) [24-28].

Discovered in 2004, HsLSD1 is a histone H3K4 mono- and di-methyl demethylase that employs flavin adenine dinucleotide (FAD) as a cofactor [29]. LSD1 has been extensively explored as a drug target due to an enzymatic activity associated with protein complexes involved in diverse biological processes [28, 30]. Indeed, dysregulation of LSD1 function has been connected to pathophysiological conditions linked with diabetes, cancer, neurodegeneration and viral infection [31, 32]. For these reasons, the development of LSD1 therapeutics has been widely investigated [33, 34]. Since the characterisation of the first LSD1 inhibitor, trans-2-phenylcyclopropylamine (2-PCPA, tranylcypromine), a large family of small molecule, mechanism-based, irreversible inhibitors that covalently modify FAD have been developed as therapeutic agents [24, 35]. Additionally, peptide analogues of the histone H3 substrate have been identified as competitive LSD1 inhibitors [25]. Additional compounds have been identified that disrupt essential protein-protein interactions within the LSD1 complex or the binding of the LSD1-containing complex to the nucleosome [36, 37].

Therefore, to build upon recent successes in small molecule targeting of SmLSD1 [19, 21] and to further identify schistosome phenotypes associated with SmLSD1 enzyme inhibition (i.e. accumulation of H3K4me2 marks), we investigated the anthelmintic activity of a library of 39 HsLSD1 inhibitors. We reveal a critical role for SmLSD1 in miracidia-to-sporocyst transitioning, schistosomula/juvenile worm survival, adult worm motility and egg production. Furthermore, we show that defects in adult worm motility and egg production are associated with reductions in neoblast and gonadal stem cell proliferation. Collectively, these data support further investigation of compounds inhibiting this histone-modifying enzyme in the pursuit of novel anti-schistosomals.

## Results

### HsLSD1 inhibitors differentially affect schistosomula phenotype and motility

A small library of LSD1 inhibitors was acquired through commercial and collaborative sources or from information derived from a recent large-scale RNAi investigation of *S. mansoni* gene function [38] (**Fig 1** and **S1 Table**). This collection included the only FDA-approved LSD1 inhibitor, trans-2-Phenylcyclopropylamine, (2-PCPA or better known as tranylcypromine (**1**) [35]), several small molecules undergoing clinical testing (including GSK-LSD1 (**2**) [39], ORY-1001 (**3**) [40] and GSK2879552 (**8**) [41]), pharmacologically active compounds synthesised around a substituted lysine scaffold (compounds **9** and **10** [42, 43], the latter currently in clinical trial for the treatment of myelofibrosis [44]) and a selection of derivatised chemicals (compounds **11**-**38**) [27, 45]. This collection also included two dual LSD1/HDAC inhibitors (compounds **6** and **7**) [26] designed as hybrid compounds resulting from the combination of a standard HDAC zinc binding group (benzamide group of Entinostat (MS-275) [46]) to either a phenelzine derivative (**4**, also known as Bizine [47]) or a cyclopropylamine analog of Bizine (**5**, [48]), respectively. This small library of LSD1 inhibitors also included a cyano-substituted indole compound (compound **39**) developed by Novartis for the treatment of LSD1-mediated diseases or disorders [49].

**Figure 1.**
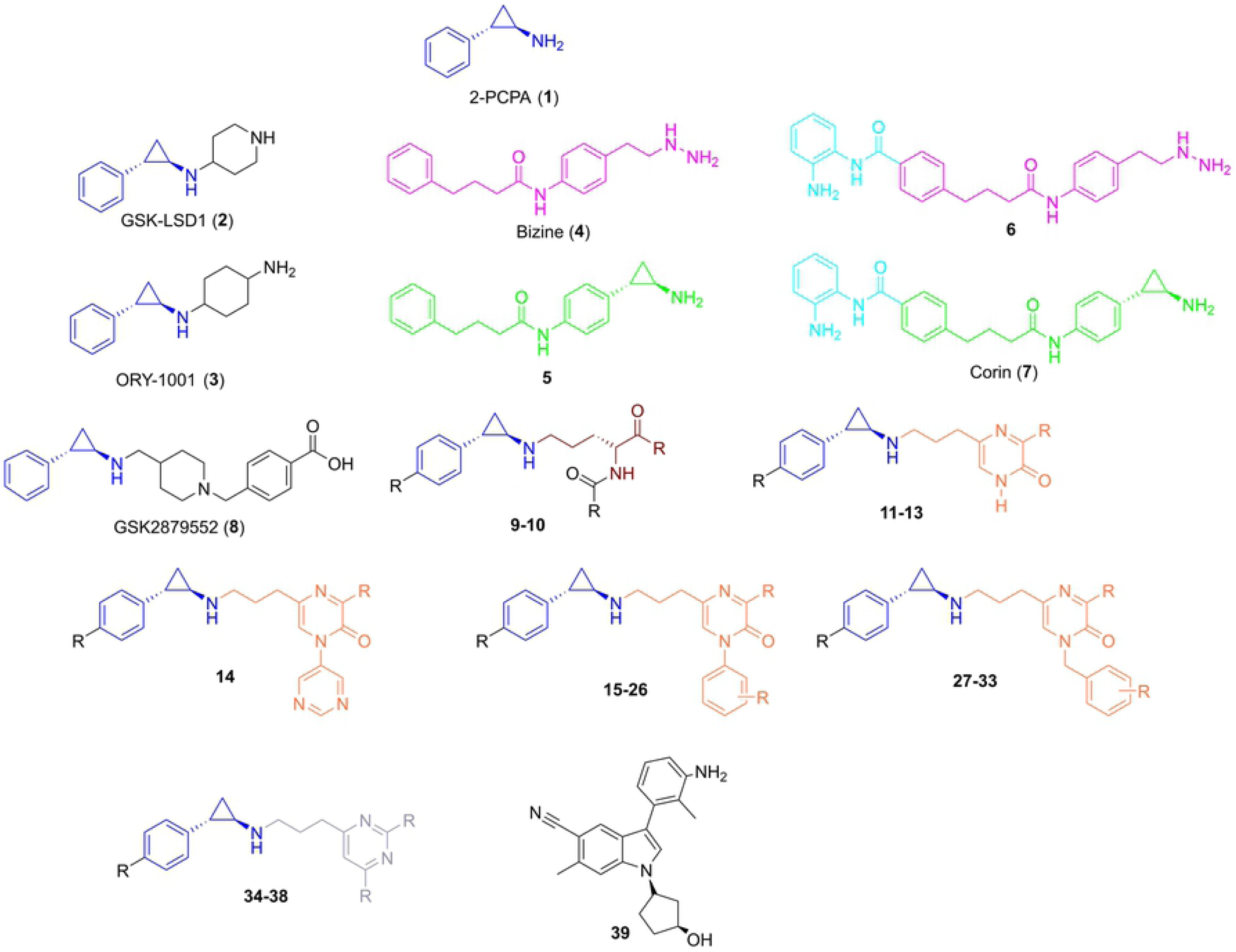
Nomenclature of LSD1 inhibitors assessed as potential anti-schistosomals. The chemical structures of tranylcypromine (2-PCPA, **1**), the first known LSD1 inhibitor, and other derived compounds developed as covalent inhibitors of LSD1 are shown. The compounds (numbered from **1** to **39**) are grouped in subclasses based on their structural similarity and a coloured code is used to highlight common structural scaffolds: blue represents the phenyl substituted tranylcypromine core; cyan indicates the incorporated features of the HDAC inhibitor (Entinostat) coupled to either the established LSD1 inhibitor bizine (in magenta) or the cyclopropylamine analogue of bizine (compound **5**, in green); brown represents the lysine mimetic scaffold; orange denotes the propylpyrazin-2-(1H)-one alone or differently substituted with pyrimidine, phenyl or benzyl cores; light blue signifies the propyl-pyrimidine scaffold. The chemical structures of each compound are reported in S1 Table. The commercial name of some compounds is also reported if known.

The selected compounds (10 µM final concentration) were initially co-cultured with schistosomula for 72 h. At this concentration, each compound was screened at least three times (biological replicates) and, in each screen, the effect of the compounds on schistosomula phenotype and motility was assessed twice (technical replicates). These whole organism assays were quantified using the in-house facility Roboworm [20, 50, 51]. For each screen, the calculated Z′ scores for both phenotype and motility metrics were within acceptable ranges (**S2 Table**) as previously described [52]. Upon screening (**Fig 2**), five compounds (**15, 16, 33, 35** and **36**, in red) were always identified as hits on both metrics (phenotype **Fig 2A** and motility **Fig 2B**) when compared to negative (0.625% DMSO) and positive (10 µM Auranofin - AUR in 0.625% DMSO) controls [53].

**Figure 2.**
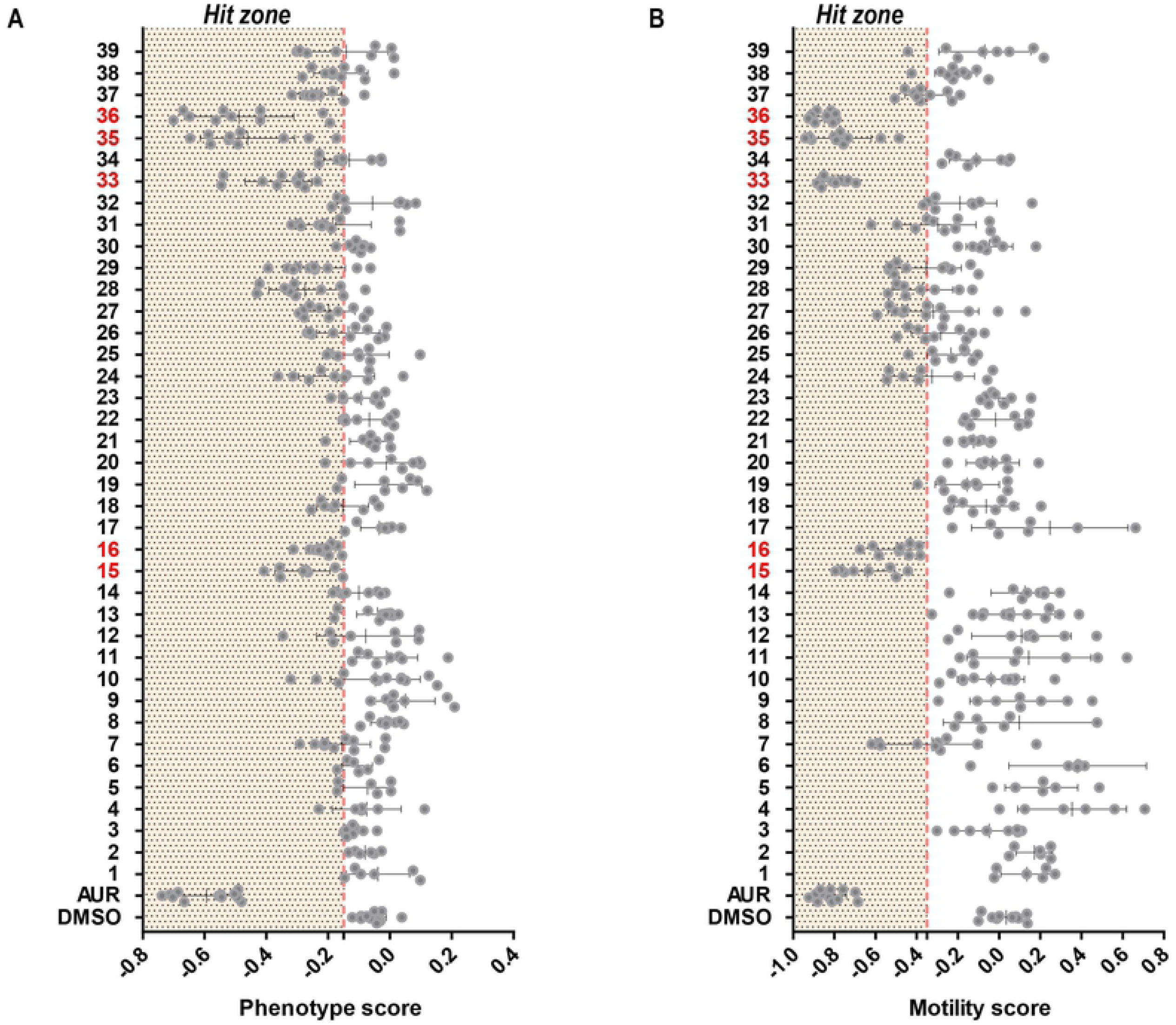
*In vitro* schistosomula screen of putative SmLSD1 inhibitors. Mechanically-transformed schistosomula (n = 120) were incubated with each of the 39 compounds for 72 h at 37°C in a humidified atmosphere containing 5% CO_2_. At 72 h, the effect that each compound had on parasite phenotype (**A**) and motility (**B**) was assessed by the high throughput platform Roboworm and compared to negative (0.625% DMSO) as well as positive (10 µM Auranofin in 0.625% DMSO) controls. Each compound was screened two to three times in three independent screens. The compound score is shown as grey dot and whiskers represent the average score and standard deviation across the screens. Compounds with activity on schistosomula phenotype and motility are shown within the ‘Hit Zone’ (delineated by the vertical dashed red lines in the graphs; - 0.15 and - 0.35 for phenotype and motility scores, respectively). All compounds showing a score lower than both reference values are considered hits (placed in an area highlighted as “Hit zone”). The five compounds highlighted in red were consistent hits across the independent screens. Z′ scores of drug screens are reported in **Table S2**.

Reassuringly, and in line with previous studies [20], GSK-LSD1 (compound **2**) failed to affect either schistosomula motility or phenotype. Amongst the five hits, compounds **33, 35** and **36** appeared to be more potent than compounds **15** and **16** (i.e., schistosomula motility and phenotype scores for the first three compounds were lower than the latter two, **Fig 2**). A subsequent dose-response titration of these five compounds confirmed this observation (Z′ scores for both phenotype and motility metrics of each screen summarised in **S3 Table**); EC_50_ values for schistosomula phenotype metrics were higher for compounds **15** and **16** (9.50 and 7.57 µM, respectively) when compared to the remaining three (4.37, 5.03 and 4.72 µM for compounds **33, 35** and **36, S1 Fig**).

### Five HsLSD1 inhibitors affect adult worm motility and IVLE production

The effect of anti-schistosomula compounds (**15, 16, 33, 35** and **36**) on adult male and female schistosome pairs (7 weeks old) was next explored to expand their anti-schistosomal applicability (**Fig 3**). Here, all compounds had a lethal effect (i.e., absence of parasite motility and gut peristalsis for 30 seconds associated with parasite detachment) on the parasite at the highest concentrations tested (50 and 25 µM, **Fig 3A**). A similar observation was recorded for all compounds at 12.50 µM; the only exception being compound **16**, which severely inhibited parasite motility, but did not lead to lethality. Upon further adult worm titrations, and consistent with the schistosomula screens, compound **33** displayed the greatest activity and inhibited schistosome motility at concentrations as low as 6.25 µM. At 3.13 μM, the effect of all compounds was minimal except for compound **33** (**S1 Movie**); at lower concentrations, the treated worms started recovering when compared to the control (**Fig 3A**).

**Figure 3.**
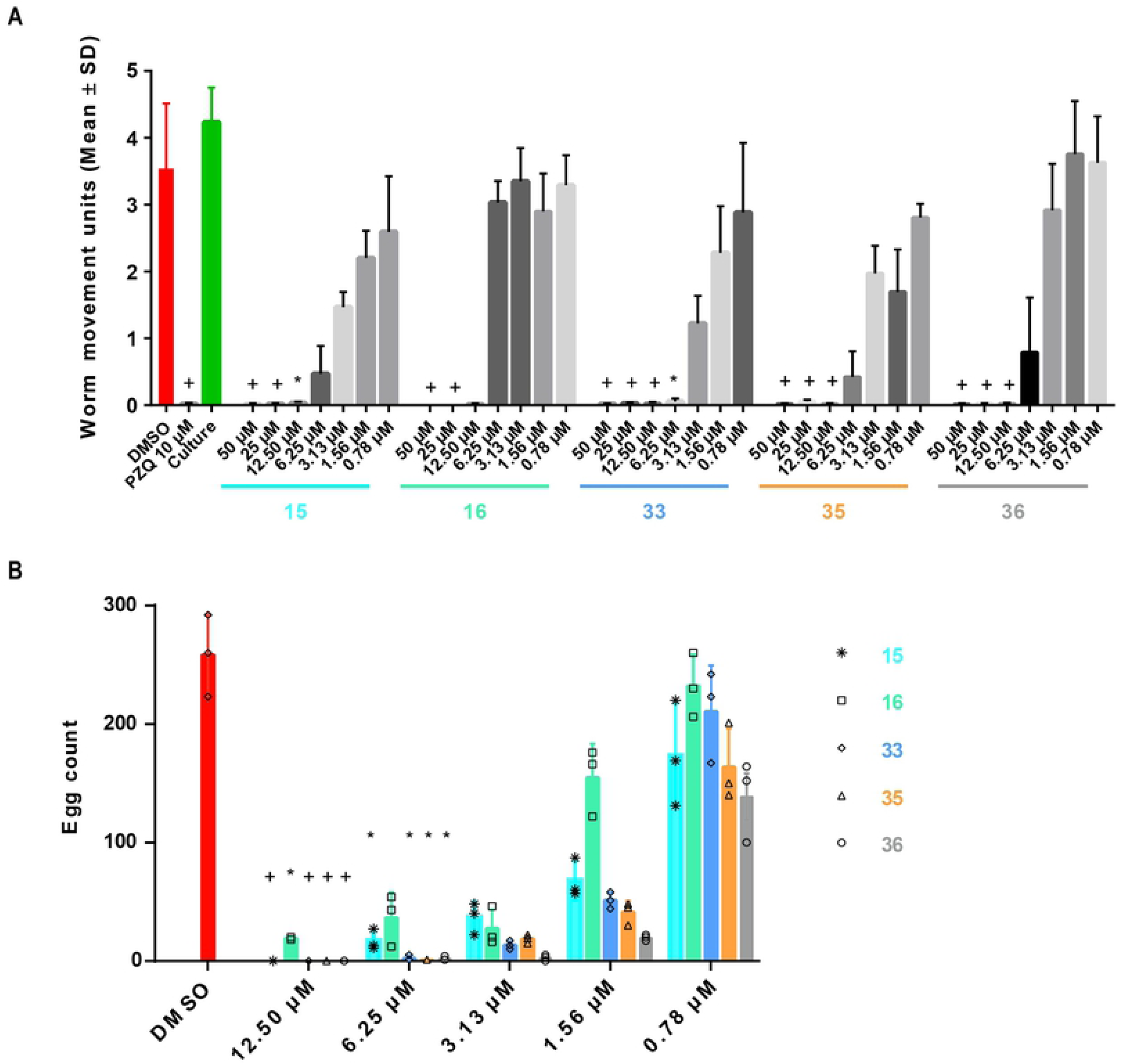
Adult worm motility and *in vitro* laid egg (IVLE) production are impaired by LSD1 inhibitors. A dose response titration (50 - 0.78 µM in 0.625% DMSO) of the five compounds was performed to assess their potency on *S. mansoni* adult worms (1 pair/well; n = 6). The titration was performed in duplicate in three independent screens. (**A**) - Worm movement was recorded at 72 h with WormassayGP2. The average worm movement (+ SD) of the three independent screens is indicated. Compound-inhibited worm movement is compared to controls (0.625% DMSO, culture control and 10 µM PZQ). (**B**) - At 72 h, eggs were collected and enumerated. For each concentration tested, individual egg counts are represented in a scatter plot; the average and standard error across the replicates is represented as a bar chart. A Kruskal-Wallis ANOVA followed by Dunn’s multiple comparisons test was performed to compare each population mean to DMSO mean. For both panels, * and + represent *p* < 0.0332 and *p* < 0.0021, respectively.

*In vitro* laid egg (IVLE) production was also affected by co-cultivation with all five compounds (**Fig. 3B**). Unsurprisingly, based on motility readouts (**Fig 3A**), no eggs were recovered from the culture media after 72 h incubation with the highest concentrations (50 and 25 µM) of all five compounds. At 12.50 µM, few eggs were recovered after co-incubation with compound **16**. However, compounds **33, 35** and **36** again demonstrated the strongest effects in inhibiting this critical process involved in host immunopathology and lifecycle transmission. For compounds **35** and **36**, IVLE production was inhibited even at a lower concentration (i.e. 6.25 µM) in which worm motility recovered. IVLE production approaching control numbers (DMSO worms) was only restored at compound concentrations of 3.13 µM. In addition to the observed reduction in egg production, a number of free (not packaged in fully formed eggs) vitelline cells, oocytes and spermatozoa were observed following treatment with a sublethal dose of compound **33** (3.13 μM, **S2 Fig** and **S1 Movie**), which was previously observed in a limited number of unrelated drug studies [54-56].

### Compound 33 inhibits vitellocyte packaging

In order to further explore the IVLE deficiencies induced by these compounds, particular attention was focused on compound **33** as representative of the five most active LSD1 inhibitors of this study (**Fig 4** and **S2 Movie**).

**Figure 4.**
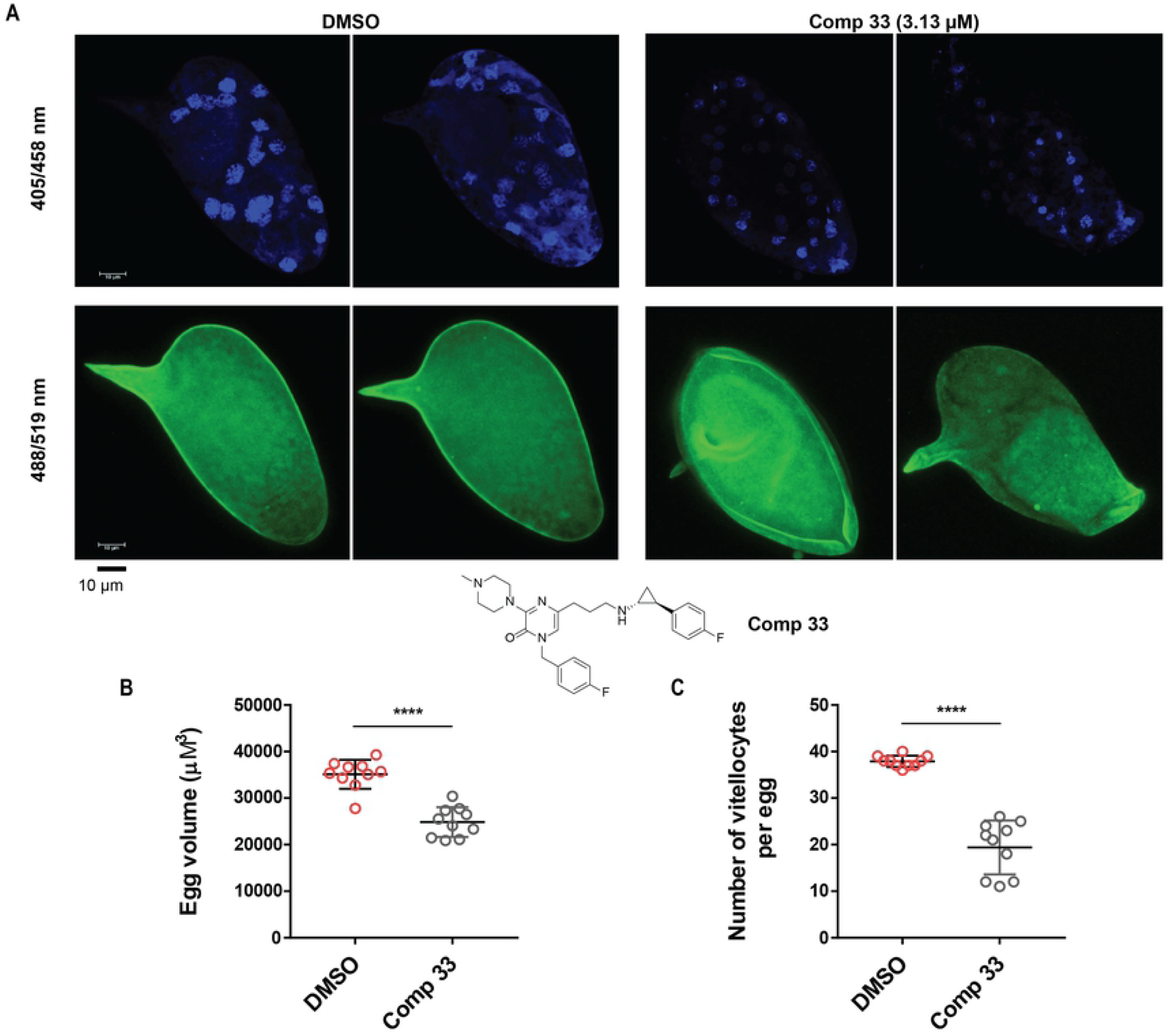
LSD1 inhibition significantly affects IVLE volume and vitellocyte packaging. Adult schistosome pairs (1 pair/well; n = 6) were co-cultured for 72 h with a sub-lethal concentration of compound **33** (3.13 μM in 0.625% DMSO) or DMSO (0.625%). Eggs were collected, enumerated and stained with DAPI prior to laser scanning confocal microscopy. (**A**) - Representative IVLE phenotypes (DAPI; Ex = 405 nm, Em = 458 nm and autofluorescence; Ex = 488 nm, Em = 519 nm) of eggs derived from the drug treated worm cultures compared to the negative control cultures (0.625% DMSO). Scale bar = 10 µm. The chemical structure of compound **33** is reported here. (**B**) - Quantification of egg volumes between treatment (compound **33**) and DMSO control groups (n = 10 per group). (**C**) - Number of vitellocytes per egg between treatment (compound **33**) and DMSO control groups (n = 10 per group). Fluorescent microscopic images (10 eggs per treatment) were acquired on a Leica TCS SP8 super resolution laser confocal microscope fitted with a 63X objective (water immersion, 1,75 zoom factor, Z stack of 60 steps) using the Leica Application Suite X. DAPI stain = blue; Autofluorescence = green. Mean and standard error are represented in Panels B (volume) and C (vitellocyte numbers). A Mann-Whitney t-test was subsequently performed to identify statistical difference between the treatments (**** corresponds to *p* < 0.0001).

Particularly, eggs derived from schistosome cultures co-incubated with sub-lethal concentrations of this chemical (3.13 µM, which did not significantly reduce worm motility - **Fig 3A**) were analysed using confocal microscopy and compared to IVLE derived from DMSO-treated worms. Even though there were no evident phenotypic abnormalities (lateral spine and oval shape were both present) in the IVLEs analysed (**Fig 4A**), a significant difference in overall egg volume was observed in compound **33** treated worms (**Fig 4B**). Moreover, following vitellocyte quantification, this compound also significantly inhibited the packaging of this critical cell population into IVLEs (**Fig 4C** and **S2 Movie**) [57, 58].

### Compound 33 inhibits adult worm stem cell proliferation and SmLSD1 activity

Due to a previous role ascribed for LSD1 in maintaining mammalian stem cell function [59], 5-ethynyl-2′-deoxyuridine (EdU) labelling of compound **33** treated adult worms (sub-lethal concentration; 3.13 µM) was performed to assess neoblast and gonadal stem cell proliferation. EdU labelling was performed on both male (**Fig 5** and **S3 Movie**) and female (**S3 Fig** and **S4 Movie**) worms treated with a sublethal concentration (3.13 µM) of compound **33** for three days.

**Figure 5.**
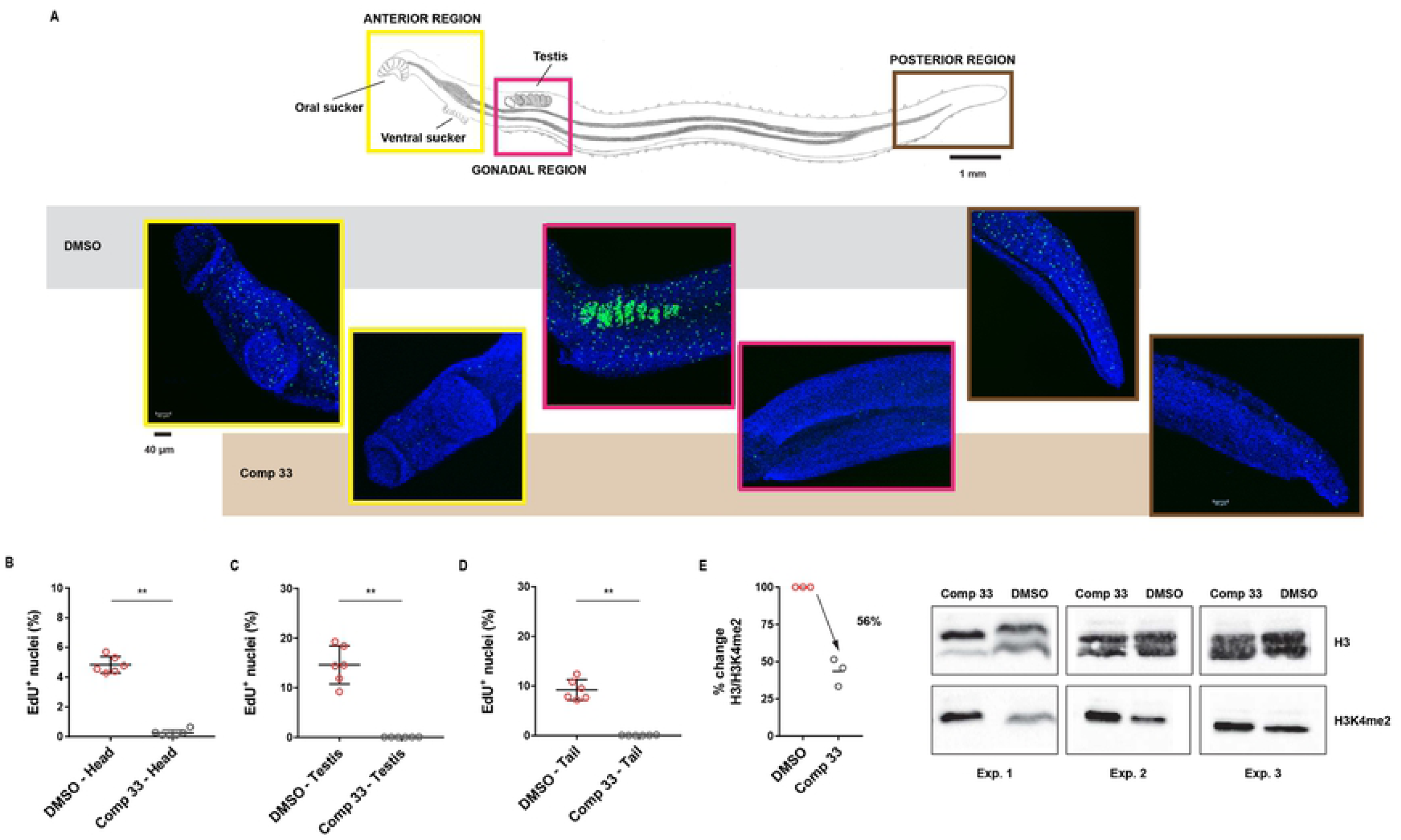
Compound 33 treatment significantly inhibits stem cell proliferation as well as H3K4me2 demethylation in adult male worms. Male schistosomes were treated with 3.13 μM of compound **33** (n = 6) or DMSO (n = 6) for 48 h and then were labelled with EdU for an additional 24 h. (**A**) - Hand drawing summarising the morphology of *S. mansoni* adult male with representative anterior region (yellow box), gonadal (magenta) and posterior region (brown) of untreated (top row, grey) compared to the drug-treated (bottom row, light brown) worms. Scatter plots illustrate the percentage of proliferative stem cells present in control (DMSO treated worms, n = 6) versus drug-treated males (3.13 µM of compound **33**, n = 6) in the head region (**B**), the testes (**C**) and the tail region (**D**). Standard errors are shown and Mann-Whitney t-test (with ** corresponding to *p* < 0.0021) was subsequently performed to quantify statistical significance between treatments. Fluorescent microscopic images (6 males per treatment) were acquired on a Leica TCS SP8 super resolution laser confocal microscope fitted with a 40X objective (water immersion, 1,00 zoom factor, Z stack of 60 steps) using the Leica Application Suite X. DAPI stain = blue; EdU+ cells = green. Scale bar represents either 1 mm or 40 μm in (**A**). (**E**) - Quantification of H3/H3K4me2 marks in adult male worm histone extracts (derived from 20 individuals, three biological replicates) after 72 h incubation with 3.13 µM of compound **33**. Western blots of each biological replicate are also reported here showing the H3 (loading controls) and H3K4me2 abundances of each experimental replicate.

For each worm (n = 6), three different anatomical regions (anterior region, gonadal system and posterior region) were observed and the number of EdU-stained dividing cells was quantified (as a percentage of all DAPI-stained cells). As shown in representative images of the parasite samples, a reduction in EdU^+^ cells was detected in drug-treated but not in control samples (both male and female parasites, **Fig 5A** and **S3A Fig**, respectively). Quantification of EdU^+^ nuclei revealed that compound **33** reduced cellular proliferation similarly across the different anatomic regions of the parasite body, regardless of sex or stem cell source (**Fig 5** and **S3 Fig**, panels **B-C-D**).

To identify if compound **33**-associated decreases in stem cell proliferation were correlated with inhibition of LSD1 demethylation activity, quantification of H3K4me2 (normalised for H3) marks in adult worm histone extracts was subsequently performed (**Fig. 5E**). Here, western blot analysis of histone extracts derived from compound **33** treated males showed an average 56% reduction in the relative ratio of H3/H3K4me2 marks (compared to the DMSO treated control) indicating specific inhibition of LSD1 activity (i.e., accumulation of H3K4me2 in the drug-treated worms (**Fig 5E**)).

### Compound 33 reduces juvenile worm viability and miracidia transformation

The activity of compound **33** was further explored on two other important *S. mansoni* life cycle stages, the immature juvenile worms and the snail-infective miracidia. Firstly, juvenile (3 weeks old) schistosomes were subjected to dose response titrations of this compound for 72 h, after which both parasite motility and viability was assessed (**Fig 6**).

**Figure 6.**
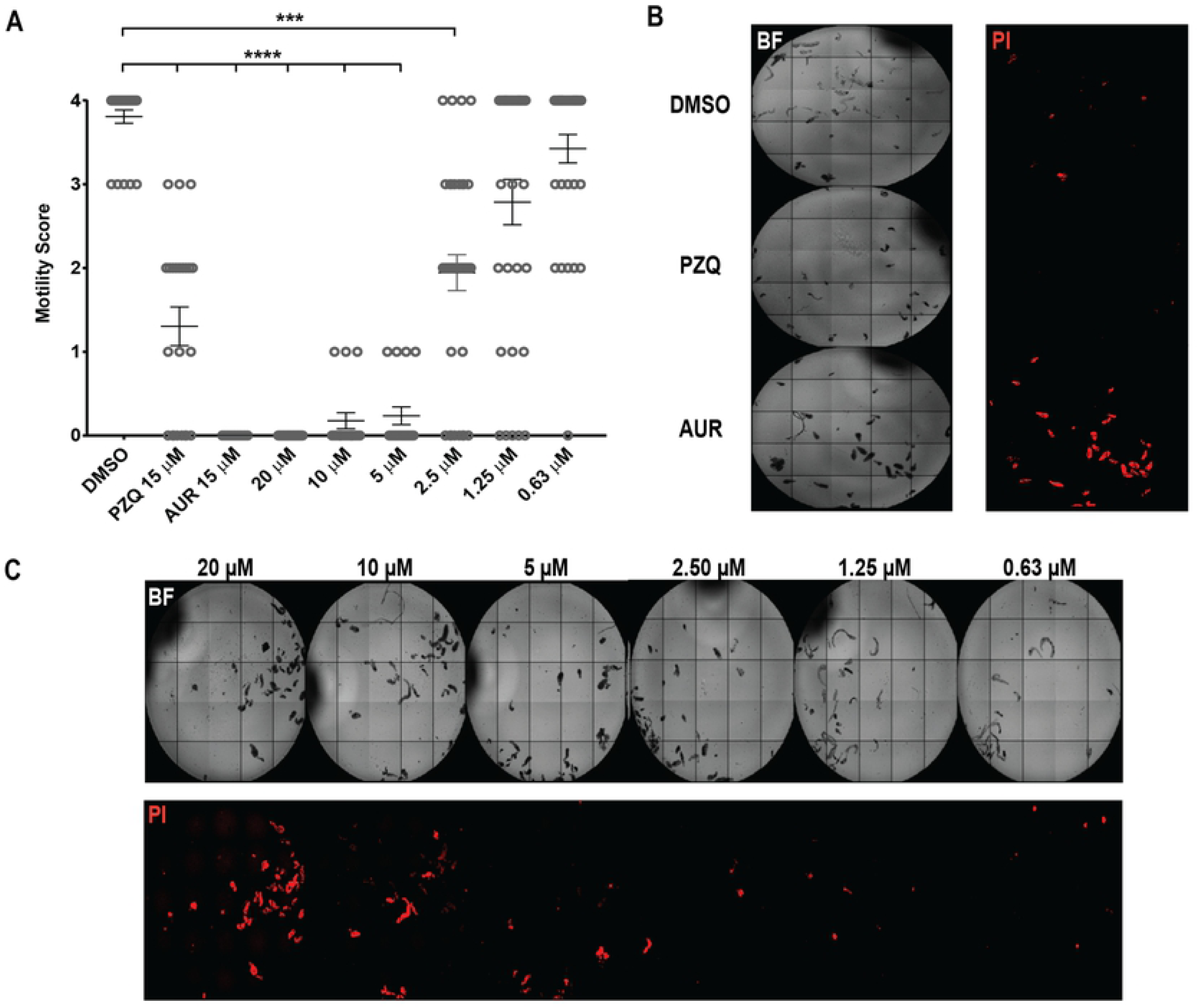
SmLSD1 inhibition leads to decreased juvenile worm motility and viability. Juvenile *S. mansoni* worms (3 weeks post infection; n = 25-30 parasites) were subjected to a dose response titration of compound **33** (20, 10, 5, 2.50, 1.25 µM and 0.63 µM in 1.25% DMSO). Motility (0 = dead, 1 = movement of the suckers only and slight contraction of the body, 2 = movement of the anterior and posterior regions only, 3 = full body movement but sluggish, 4 = normal movement) and viability metrics (PI positive parasites) were assessed at 72 h post-dosing and compared to control parasites (negative control: 25-30 juveniles co-cultivated in the presence of 1.25% DMSO; positive controls: 25-30 juveniles co-cultured in either 15 µM PZQ or AUR in 1.25% DMSO). (**A**) - The scatter plot shows the motility score for each parasite/treatment. The mean motility and the standard error of the mean motility are also included in the plot. A Kruskal-Wallis ANOVA followed by Dunn’s multiple comparisons test was performed to compare each population mean to DMSO mean. ***, **** represent *p* < 0.0002, *p* < 0.0001, respectively. (**B**) - Representative images of PI-stained (2 µg/ml) juveniles treated with DMSO, praziquantel (PZQ, 15 µM) and auranofin (AUR, 15 µM). (**C**) - Representative images of compound **33**/parasite co-cultures showing a concentration-dependent increase in PI staining. The plate was imaged under both bright-field (BF) and fluorescent (for propidium iodide detection, PI) settings, using an ImageXpressXL high content imager (Molecular Devices, UK).

At the highest concentration tested (20 µM), compound **33** significantly reduced parasite movement when compared to the negative (DMSO) control (**Fig 6A**). A similar observation was recorded for AUR-treated parasites (at 15 µM). Furthermore, when visualised for propidium iodide (PI) uptake at 536 nm (**Figs 6B** and **6C**), both treatments were associated with increased fluorescence, providing confirmation of juvenile worm death. In line with other reports [6, 60, 61], PZQ only showed partial activity on juvenile worms (**Fig 6A**), and as confirmed by a lower uptake of PI (**Fig 6B**), these parasites where not all dead. At lower concentrations of compound **33** (10, 5 and 2.50 µM), reductions in both motility (**Fig 6A**) and viability (**Fig 6C**) were still noted when compared to the DMSO control. However, at lower concentrations (1.25 µM and 0.63 µM), juveniles started recovering normal movement and mortality was reduced.

Although drug screens were mainly focused on intra-mammalian parasitic stages, we were also interested in whether the most potent LSD1 inhibitor affected schistosome developmental forms that interact with the intermediate molluscan host [62]. Hence, free-swimming miracidia were exposed to compound **33** (in a dose response titration) for 48 h and *in vitro* transformation into sporocysts was assessed (**Fig 7**).

**Figure 7.**
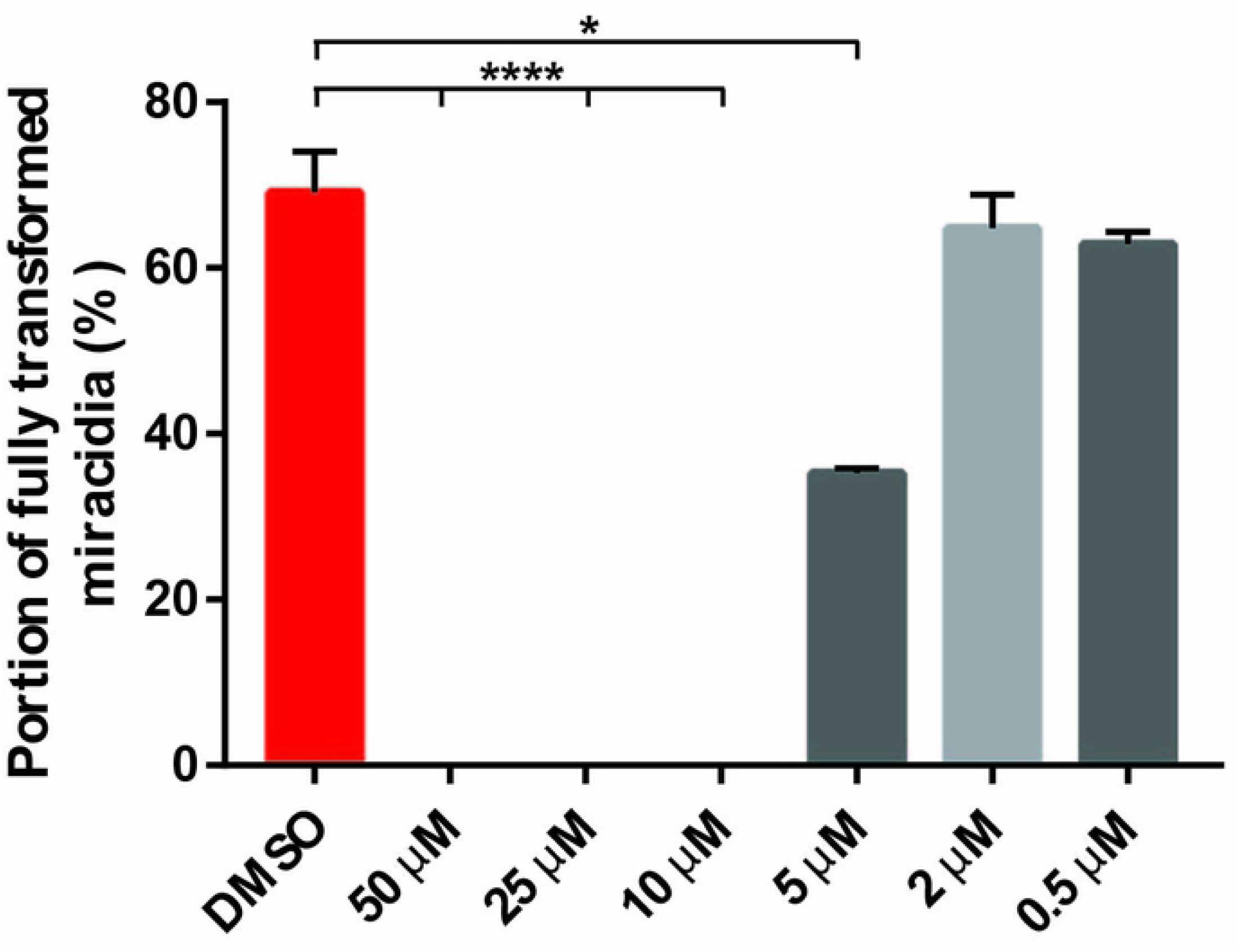
SmLSD1 inhibition blocks miracidia to sporocyst transformation. Miracidia were exposed to compound **33** during a dose response titration (50, 25, 10, 5, 2 and 0.5 µM in 1% DMSO). Sporocyst transformation was scored (%) after 48 h. Each titration point was conducted in triplicate and compared to parasites cultured in CBSS with 1% DMSO (controls) at a constant temperature of 26°C (in the dark). Means and standard error are shown. A Kruskal-Wallis ANOVA followed by Dunn’s multiple comparisons test was performed to compare each population mean to DMSO mean. * and **** represent *p* < 0.0332 and *p* < 0.0001, respectively.

Upon titration, a significant inhibition in miracidia to sporocyst transformation was found for parasites treated with 50 - 5 µM of compound **33**. In fact, no movement or flame cell activity (related to parasite death) was observed at 50, 25 and 10 µM concentrations. At 5 µM, compound **33** inhibited miracidia-sporocyst transformation by 66%. Below 5 µM, miracidia to sporocyst transformation was unaffected.

### The five selected compounds exhibit different parasite-host selectivity

Overt cytotoxic activity was next explored on human HepG2 cells by prioritising those compounds that showed the most potent anti-schistosomal effects (compound **15, 16, 33, 35** and **36**). A previous large scale mammalian cytotoxicity study indicated that maximal HepG2 cytotoxicity was observed within the first 24 h for 91% of the active compounds [63]; therefore, 24 h continuous co-incubation of compounds with HepG2 cells was selected for this study to provide preliminary evidence of overt cytotoxicity. Each compound was tested in a dose response titration (200 to 0.01 µM, **S4 Fig**), with the average CC_50_ of each compound reported in **Table 1**.

**Table 1.** Anti-schistosomal activity and HepG2 cytotoxicity summary of the most potent LSD1 inhibitors.

| Compound | $\text{EC}_{50}$<br>(schistosomula, P, $\mu\text{M}$) | $\text{EC}_{50}$<br>(adults, $\mu\text{M}$) | $\text{CC}_{50}^*$<br>($\mu\text{M}$) |
| --- | --- | --- | --- |
| **15** | 9.503<br>(7.537-12.09)<sup>a</sup> | 2.431<br>(1.841-3.148)<sup>a</sup> | 28.4<br>(22.52-34.44)<sup>a</sup> |
| **16** | 7.569<br>(4.062-13.49)<sup>a</sup> | 7.330<br>(6.373-9.234)<sup>a</sup> | 13.54<br>(11.84-15.14)<sup>a</sup> |
| **33** | 4.370<br>(3.412-5.764)<sup>a</sup> | 2.137<br>(1.714-2.634)<sup>a</sup> | 2.765<br>(1.679-3.743)<sup>a</sup> |
| **35** | 5.031<br>(3.985 - 6.346)<sup>a</sup> | 2.293<br>(1.742-2.965)<sup>a</sup> | 20.82<br>(18.81-23.02)<sup>a</sup> |
| **36** | 4.719<br>(3.869 - 5.872)<sup>a</sup> | 4.560<br>(3.697-5.519)<sup>a</sup> | 26.08<br>(21.62-32.44)<sup>a</sup> |

Schistosomula $\text{EC}_{50}$ (derived from phenotype metrics, P) values of the compounds are calculated based on three dose response titrations (10 - 0.625 $\mu\text{M}$ ). Adult worm $\text{EC}_{50}$ values of the compounds were calculated based on three dose response titrations (50 - 0.78 $\mu\text{M}$ ). Parasite screens were conducted for 72 h.
\* The $\text{CC}_{50}$ values were calculated based on three dose response titrations (200 - 0.01 $\mu\text{M}$ ) using a HepG2 cell line co-cultivated with the compounds for 20 h (followed by another 4 h with the MTT reagent) [63]. <sup>a</sup> 95% confidence interval.

Four of the five compounds displayed intermediate levels of HepG2 cytotoxicity with CC_50_s higher than 13 µM (compounds **15, 16, 35** and **36**). Compound **33**, the most active LSD1 inhibitor on both larva and adult stages, also appeared to be the most cytotoxic (CC_50_ below 10 µM, **Table 1**).

### Compound 33 likely inhibits SmLSD1 activity by covalently reacting with the FAD cofactor

Irreversible inhibitors of LSD1 (tranylcypromine and its derivatives) have been shown to modify the FAD cofactor by covalent bonding of their cyclopropylamine group to the N5 atom of the cofactor flavin ring. Hence, the resulting N5 adduct inhibits the demethylase activity of the enzyme [24]. Due to structural similarity of compound **33** with these tranylcypromine derivatives [64-66], a similar molecular mechanism of action was expected upon interaction of this compound with SmLSD1. To explore this, homology modelling and compound docking were performed (**Fig 8**).

**Figure 8.**
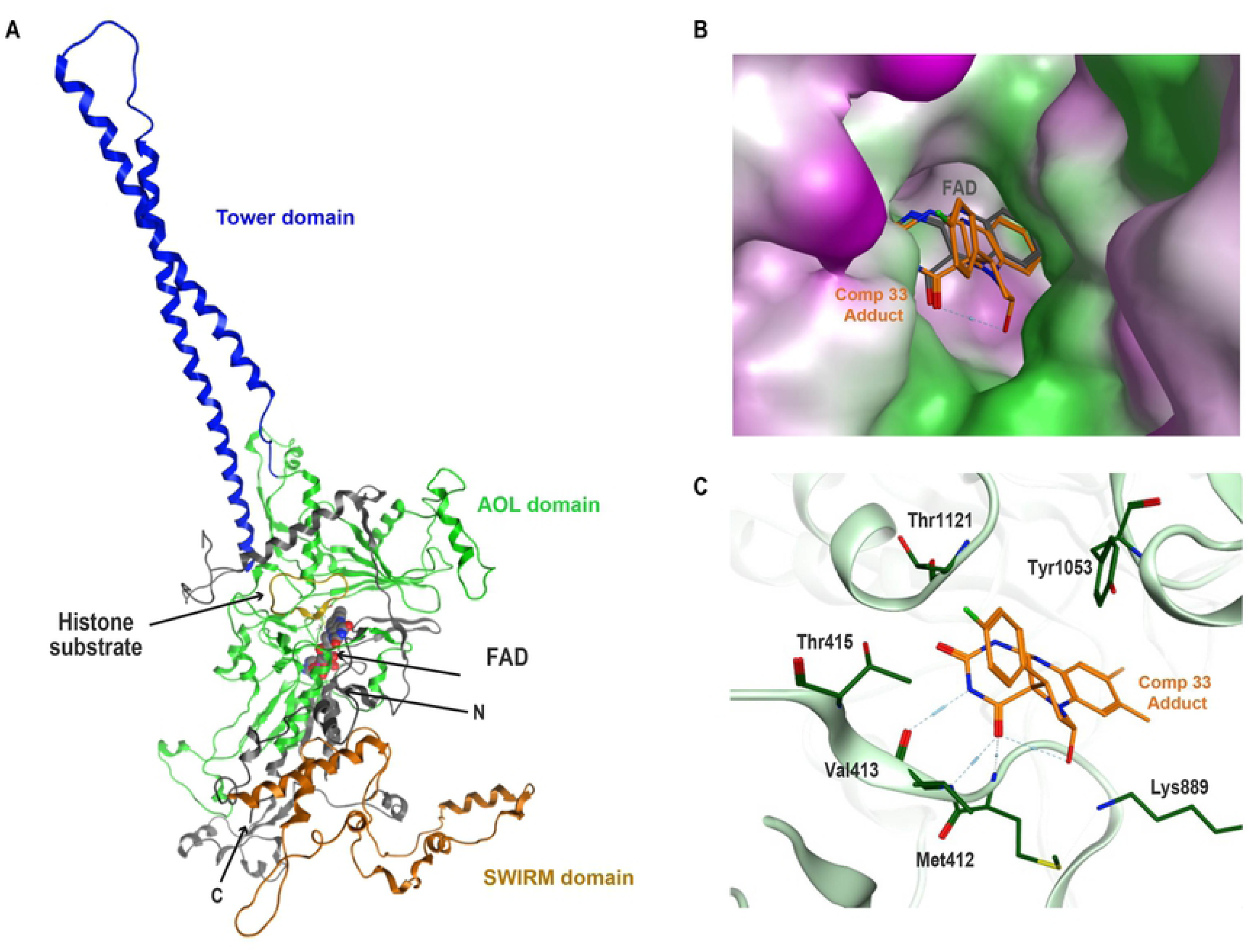
Proposed binding mode of the compound 33-derived adduct in SmLSD’s active site. (**A**) - Ribbon diagram representation of full-length Smp_150560’s homology model: unstructured N-terminal region (N) in grey, SWIRM domain in orange, AOL domain in green and Tower domain in blue. The C-terminus is indicated (C); the cofactor FAD is shown as spheres (grey for carbons, red for oxygen, blue for nitrogen); the histone protein is represented as yellow ribbon; the methionine is shown as yellow stick. (**B**) - Surface diagram of SmLSD1 showing the FAD−PCPA adduct of compound **33** in the protein’s active site. The binding mode of this adduct is compared to the orientation of the cofactor FAD (here the structure of the cofactor was reduced at only the flavin ring). The adduct and the flavin ring of FAD are shown as a stick model in orange and dark grey, respectively. The protein surface was coloured by lipophilicity with purple, white and green representing hydrophilic, neutral and lipophilic regions, respectively. (**C**) - Ligand interactions of the compound **33** covalent adduct with highlighted conserved active site residues of SmLSD1. Amino acid lateral chains involved in interactions are shown as dark green sticks and labelled according to their position in the full-length amino acid sequence of Smp_150560 (shown as light green ribbon).

As previously published [19], the homology model of the parasite enzyme was generated using HsLSD1 (PDB entry: 2V1D, **Fig 8A**) [67]. In order to predict the binding mode of compound **33** (**S5A Fig**) to SmLSD1, the N5 adduct of this compound with the flavin ring of the FAD (**S5B Fig**) cofactor was prepared (**S5C Fig**) and used as a ligand for molecular docking to the SmLSD1 homology model. The predicted docking pose fitted very well in the active site of SmLSD1 with the flavin ring of the adduct superimposed on the same tricyclic ring of the FAD cofactor (**Fig 8B**). A closer analysis of the ligand-protein interactions (**Fig 8C**) revealed many interactions between the flavin ring of the chemical adduct and amino acid residues, which are typically involved in the cofactor binding of the N-terminal region of the amino-oxidase like domain of LSD1 (Met412, Val413 and Thr415). The fluoro-phenyl ring of the FAD-adduct extended outside the active site, towards the substrate binding pocket and was embedded between two residues (Tyr1053 and Thr1121, **Fig 8C**). These two residues together define the aromatic cage, which has been observed in other amine oxidase enzymes (including LSD1) and contributes to the active site hydrophobic shielding from the influx of external solvents [68, 69]. In addition to this, we also observed the orientation of the compound towards another conserved residue, the invariant lysine (Lys889), which has been investigated for its role in catalysis as well as in proton transfer (acting as an active-site base of LSD1) [70, 71].

## Discussion

Studies of the *S. mansoni* lifecycle have shown that transitions between, and development within, both intermediate and definitive hosts are processes carefully regulated by epigenetic factors [17, 18, 23, 72-74]. While a critical role for histone methylation and demethylation in these processes has been demonstrated [19-21, 23], the specific contributions of enzymes catalysing these reversible post-translational modifications have not been thoroughly characterised. Following on from our previous investigations demonstrating that anti-neoplastic anthracyclines could bind to the target pocket of SmLSD1 and kill multiple schistosome lifecycle stages [19], we pursued the evaluation of 39 HsLSD1 inhibitors as potentially more potent anti-schistosomals.

Initial *in vitro* schistosomula screening of the 39 compounds identified five (compounds **15, 16, 33, 35** and **36**) with activity (EC_50_ below 10 µM; **Fig 2** and **Table 1**) similar to that previously found for the putative SmLSD1 inhibitors daunorubicin hydrochloride (EC_50_ below 6 µM) and pirarubicin (EC_50_ below 3 µM) [19]. When the *in vitro* screens were expanded to adult parasites (**Fig 3**), the most active anti-schistosomula compound (in this case compound **33**) also demonstrated the strongest effects (decreased worm motility and IVLE production; **Fig 3**). While a reduction in adult worm viability and IVLE production was also observed for a recently described LSD1 inhibitor (MC3935) [21], the activity of compound **33** reported here was more potent.

To potentially help explain potency differences of the 39 tested compounds, physiochemical properties such as molecular weight (MW) and lipophilicity (in terms of calculated LogP) were compared (**S6 Fig**). Most of the compounds have a molecular weight between 400 and 600 g/mol, but variable lipophilicities. These two properties do not adequately explain differences in activity among the five most potent compounds or between them and the remaining less active ones.

The presence a para-fluorophenyl substitution on the cyclopropyl ring of the tranylcypromine derivatives was identified as a common feature of the five most active compounds (**S4 Table**). Considering the mechanism of action of these compounds, they are all capable of forming a covalent adduct with FAD that contains the fluorophenyl ring. Though the para-fluro substitution may be important for potency, this structural feature cannot explain the differences in activity among the five compounds. However, for a more detailed structure-activity relationship (SAR) study, two subfamilies can be identified; the first subfamily (including compounds **15, 16** and **33**) contains an alkyl linker between the cyclopropyl ring and a pyrazine ring which is replaced by a pyrimidine ring in the second subfamily (including compounds **35** and **36**). The *in silico* molecular docking of these five compounds to SmLSD1’s active site highlighted a more favourable docking score (XP score, **S4 Table**) for compound **33;** this is likely due to the presence of a benzyl group conferring more flexibility when compared to the other compounds. This structural feature could explain the greater activity of compound **33** on both schistosomula and adult worms (**Table 1** and **S4 Table**).

Reassuringly, compound **33**-induced inhibition of IVLE production broadly recapitulated the previously-described RNAi-mediated knock down of *smlsd1* (Smp_150560) in adult worms and the viability assays of other synthetic LSD1 inhibitors [19, 21]. In addition to an egg laying deficit, compound **33** significantly reduced the overall volume of IVLEs as well as the packaging of fertilised ova/vitellocytes into them (**Fig 4** and **S2 Fig**). These collective phenotypes pointed to a global defect in the female’s egg laying machinery (ootype, vitellaria and ovary), which was further supported by compound **33**’s complete inhibition of gonadal and vitellaria stem cell (S1) proliferation (**S3 Fig**). The additional inhibition of gonadal stem cell proliferation in the testes (**Fig 5**) following drug treatment suggests an essential contribution of SmLSD1 in both female and male germline tissues (leading to defects in oogenesis, vitellogenesis and spermatogenesis). When examined further, these phenotypes appear dependent on SmLSD1 demethylase activity as H3K4me2 marks remained abundant in compound **33** treated males (**Fig 5E**). In support of this contention, it was previously demonstrated that global accumulation of H3K4me2 in *spr 5* (*lsd1* homologue) *C. elegans* null mutants led to progressive sterility in progeny due to dis-regulation of spermatogonia-associated genes [75]. When taken together, our data supports previous observations [16] and highlights an essential role for post-translational histone modifications in the control of oviposition.

In addition to affecting schistosomula and adult worm phenotypes, we also demonstrated that compound **33** markedly reduced the transformation of miracidia into germinal cell-enriched sporocysts (**Fig 7**) [76]. A ‘block in transformation’ phenotype was also observed when the histone methyltransferase inhibitors A366 and GSK343 (likely targeting G9a/GLP and EZH1/H2 homologs respectively [23]) were used as part of studies investigating the role of H3K27 methylation during schistosome lifecycle progression [23]. These combined results, derived from distinct studies of different histone methylation and demethylation components (G9a/GLP and EZH1/H2 HMTs [23] vs LSD1 HDM here), mutually support a critical role for histone methylation regulation (on both H3K4 and H3K27) in the development of miracidia to sporocyst. When further considering the activity of compound **33** on juvenile worms (**Fig 6**), collectively, our data suggests that targeting SmLSD1 with small molecules represents a strategy capable of affecting many (if not all) schistosome lifecycle stages. If compound selectivity for SmLSD1 (limiting mammalian cell cytotoxicity) could be improved (**Table 1**) through medicinal chemistry optimisation, treating schistosomiasis with SmLSD1 inhibitors offers a distinct advantage over PZQ (ineffective against all schistosome lifecycle stages [5, 77]). In fact, LSD1 inhibitors currently in clinical development for the treatment of hematologic malignancies and solid tumours have been shown to be relatively safe and well tolerated [42], suggesting that LSD1 inhibitors (or derived compounds) may be as well or better tolerated in patients with schistosomiasis.

Similar to its effect on germinal stem cells, compound **33** also inhibited adult neoblast proliferation in both sexes (**Fig 5** and **S3 Fig**). These proliferation defects are comparable to that observed in *Hslsd1*-expressing neural stem cells treated with the LSD1 inhibitors pargyline or tranylcypromine [59] and support previous findings confirming that *Smlsd1* is also expressed in rapidly dividing cells throughout the parasite [78]. *Smmbd2/3* (encoding an epigenetic reader of 5-methylcytosine) and *Smcbx* (encoding an epigenetic reader of methyl lysine) are also co-expressed in proliferative schistosome cells (*h2b+*); similar to SmLSD1 inhibition, knockdown of either reader results in reduced neoblast proliferation [79]. When additionally considered alongside neoblast proliferation defects found in adult schistosomes treated with 5-azacytidine (a DNA methyltransferase inhibitor [15, 80]), epigenetic processes are rapidly emerging as essential regulators of schistosome stem cell biology.

In conclusion, these findings demonstrate the rationale for exploring compounds developed against human targets for use as anti-schistosomals that inhibit essential functions of the parasite, and, more generally, support this approach for other NTDs caused by parasitic worms [81]. This work, alongside others [19, 21], validates SmLSD1 as a target for the treatment of disease caused by *S. mansoni*. Further optimisation of compound **33** should result in the identification of more potent and selective SmLSD1 inhibitors (more active against the parasite target than the host) facilitating the progression of urgently needed, next-generation chemotherapies for this disease.

## Experimental procedures

### Ethics statement

All procedures performed on mice adhered to the United Kingdom Home Office Animals (Scientific Procedures) Act of 1986 (project license P3B8C46FD) as well as the European Union Animals Directive 2010/63/EU and were approved by Aberystwyth University’s Animal Welfare and Ethical Review Body (AWERB).

### Compound preparation and storage

All compounds were received as dry powder. Upon delivery, each compound was dissolved in dimethyl sulfoxide (DMSO) at a stock concentration of 10 mM and a working concentration of 1.6 mM (or lower, if needed). Both stock and working solutions were stored at −20°C prior to use. Positive controls for *S. mansoni* screens included praziquantel (PZQ, P4668, Sigma-Aldrich, UK) and auranofin (AUR, A6733, Sigma-Aldrich, UK); these were also solubilised in DMSO and stored as described above.

### Ligand preparation, protein preparation and molecular docking

From the chemical structure of compound **33** (an N-alkylated tranylcypromine derivative) and the cofactor FAD (**S5A Fig** and **S5B Fig**, respectively), the covalent adduct was generated using the builder tool in the software MOE (Molecular Operating Environment) 2015.10 [82]. In summary, the chemical structure of FAD was firstly obtained from a selected crystal structure of HsLSD1 (PDB entry: 6NQU, 41 % sequence identity) and then its structure was simplified since only the flavin ring of the cofactor is involved in the mechanism of action of this class of LSD1 irreversible inhibitors. Thereafter, the desired ligand was obtained by forming a covalent adduct between the amino group of the cyclopropyl core of compound **33** and the flavin ring of the cofactor. As a result, the substitution on the amino moiety of this trans-2-phencylcyclopropylamine analogue (which most likely acts as lysine fragment mimic) is lost as shown in **S5C Fig**.

The obtained N5 adduct was saved in a *sdf* format prior to processing by the Lig Prep tool within Maestro v10.1 [83]. Twenty-five conformers of the ligand were generated and used for docking simulations. The homology model of Smp_150560 was generated within the MOE 2015.10 homology tool using a single template approach as previously described [19]. Similarly to ligand preparation, the structure of the cofactor (used for the induced-fit homology modelling of SmLSD1) was simplified. Here, only the tricyclic ring of FAD was kept whereas the 5′-adenosyldiphosphoribityl group at position 10 of the flavin ring (oriented towards the interior of the protein) was removed using the builder tool in MOE. The generated model was prepared with the Schrodinger Protein Preparation Wizard tool using the OPLS_2005 force field where hydrogens atoms were added, partial charges were assigned and energy minimisation was performed. To facilitate docking, the flavin ring of FAD was selected as the centroid to computationally prepare a 12 Å docking grid. Docking simulations were performed using the Glide docking software and the in-built Extra Precision (XP) scoring function in order to estimate the target-compound binding affinity (as expressed as XP score). Selecting default parameters, only ten output poses (conformations) for ligand conformer were generated in the final step.

### Parasite maintenance and preparation

The NMRI (National Medical Research Institute) Puerto Rican strain (PR-1) of *S. mansoni* was used to maintain the life cycle. The mammalian-specific aspect of the life cycle was passaged through *Mus musculus* (HsdOLa:TO - Tuck Ordinary; Envigo, UK); the molluscan aspect of the life cycle was maintained through two *Biomphalaria glabrata* strains - the NMRI albino and pigmented outbred strains [84].

*S. mansoni* cercariae were obtained from infected *B. glabrata* snails after 1 h of incubation at 26°C under intensified lighting conditions. Cercariae were collected in falcon tubes and incubated on ice for at least 1 h prior to transformation. Cercariae were then mechanically transformed into schistosomula as previously described [85].

*S. mansoni* juvenile or adult worms were recovered by hepatic portal vein perfusion [86] of mice previously infected for 3 weeks with 4,000 cercariae/mouse or 7 weeks with 180 cercariae/mouse, respectively. Following perfusion, juvenile worms were collected by gravity in a 50 ml falcon tube, re-suspended in clear DMEM and washed three times (300 x *g* for 2 min) to remove residual host contamination. Adult worms were separated from perfusion media by sedimentation and subsequently washed a further three times in pre-warmed adult worm media (DMEM (Gibco, Paisley, UK) supplemented with 10% (v/v) FCS (Gibco, Paisley, UK), 1% (v/v) L-glutamine (Gibco, Paisley, UK) and an antibiotic mixture (150 Units/ml penicillin and 150 µg/ml streptomycin; Gibco, UK)). All parasite material was subsequently transferred into a petri dish and incubated in a humidified environment containing 5% CO_2_ at 37°C for at least one hour. Before downstream manipulation, any macro residual host material (e.g. mouse hair, blood clots) was removed using a clean paintbrush.

To obtain miracidia, parasite eggs were isolated from 7 weeks infected mouse livers and exposed to light to induce miracidia hatching in 1X Lepple water [23, 87]. Following hatching of miracidia, parasites were incubated on ice for 15 min and then centrifuged at 700 x *g* for 5 min at 4°C. The miracidia pellet was then re-suspended in 5 ml of Chernin’s balanced salt solution (CBSS) [23], subjected to pelleting and two subsequent washes (all at 700 x *g* for 5 min at 4°C). Afterwards, the supernatant was carefully removed with a serological pipette and the miracidia-enriched pellet was resuspended with CBSS supplemented with 500 µl of Penicillin-Streptomycin (containing 10,000 units penicillin and 10 mg streptomycin/ml, P4333, Sigma-Aldrich).

### *In vitro* schistosomula screens

Schistosomula drug screens were performed using an in-house facility, Roboworm, which standardised assessment of larva motility and phenotype [51]. As previously described [50, 51], each compound (as single concentration or two-fold titrations) was transferred into individual wells of a 384-well tissue culture plate (Perkin Elmer, cat 6007460) prior to addition of 120 parasites. Each plate contained negative (0.625% DMSO) and positive (AUR at 10 µM final concentration in 0.625% DMSO) control wells. Schistosomula/compound co-cultures were then incubated at 37°C for 72 h in a humidified atmosphere containing 5% CO_2_. At 72 h, tissue culture plates were imaged under the same conditions (37°C for 72 h in a humidified atmosphere containing 5% CO_2_) using an ImageXpressXL high content imager (Molecular Devices, UK) with subsequent images processed for phenotype and motility as previously reported [51].

The phenotype and motility scores were used to evaluate whether a compound displayed anti-schistosomula activity; here, −0.15 and −0.35 defined threshold anti-schistosomula values for phenotype and motility scores, respectively. The Z′ value, a metric used to evaluate the success of high throughout screens by comparing means and standard deviations of positive and negative controls [52], of each plate was subsequently calculated. Plates with a Z′ value below the value of 0.3 were considered failed and the screen was repeated.

### *In vitro* juvenile worm screens

Twenty-five to thirty (25-30) juveniles were added to wells of a clear 96 well, flat-bottom plate and co-incubated with compounds (compound **33**: 20 - 0.63 µM in 1.25% DMSO; negative control: 1.25% DMSO; positive controls: 15 µM PZQ or 15 µM AUR in 1.25% DMSO; media only) in 200 µl of adult worm medium as previously described [50]. Treated juvenile worms were incubated for 72 h in a humidified environment containing 5% CO_2_ at 37°C. After 72 h, an adapted version of the WHO-TDR scoring system [88] was used to quantify the effect of the drug on both phenotype and motility of the parasite [50]. Juvenile worm viability was additionally quantified as previously reported with minor modifications [89]. Briefly, propidium iodide (PI, P1304MP, Sigma-Aldrich, UK) was added to juvenile/compound co-cultures (to a final concentration of 2 µg/ml) and these co-cultures were imaged under both bright-field and fluorescent settings (PI detection, 535 and 617 excitation and emission wavelengths, respectively), using an ImageXpressXL high content imager (Molecular Devices, UK).

### *In vitro* adult worm screens

Adult worms (1 worm pair/1 ml of adult worm media) were dosed with compounds at final concentrations spanning 50 µM - 0.78 µM (in 0.5% DMSO) in 48 well tissue culture plates. DMSO (0.5%) and praziquantel (10 µM in 0.5% DMSO) were also included as negative and positive control treatments. Treated adult worms were incubated for 72 h in a humidified environment at 5% CO_2,_ 37°C. Parasite motility after drug treatment was assessed by a digital image processing-based system (WormassayGP2) [38, 90] modified after Wormassay [91, 92].

Where egg deposition was noticed, eggs were removed and counted using a Sedgewick rafter. After counting, eggs were immediately transferred to a 1 ml microfuge tube and centrifuged at 200 x *g* for 2 min. With the eggs loosely pelleted at the bottom of the microfuge tube, the remaining media was carefully removed, and the egg pellet was fixed in formalin (10% v/v formaldehyde).

### *In vitro* miracidia screens

Miracidia (20-50 individuals in CBSS) were transferred to a 24-well tissue culture plate and dosed with a titration of compound **33** (50, 25, 10, 5, 2 and 0.5 µM in 1% DMSO) [23]. Each treatment was performed in duplicate; parasites cultured in CBSS with 1% DMSO were included as a negative control. Miracidia were incubated for 48 h at 26°C and subsequently evaluated for morphological and behavioural changes differing from control cultures (1% DMSO) using an Olympus inverted light microscope. Dead, fully transformed and partially transformed miracidia were enumerated in the DMSO control and the drug cultures as previously described in literature [12, 23, 62]. Miracidia were scored as fully transformed if the transformation process was completed, the sporocyst surface was fully formed, no cilia plates remained attached and normal movement was detectable. A miracidium was scored as partially transformed if the parasite was no longer swimming, assumed a rounded morphology and ciliated plates remained attached to the parasite surface. Dead parasites were identified if they did not show any signs of movement, extensive degradation was present at the surface and no flame-cell activity was evident.

### Cytotoxicity assay

Human Caucasian Hepatocyte Carcinoma (HepG2) cells (85011430, Sigma Aldrich, UK) were used to assess the overt cytotoxicity of selected compounds. A MTT (3-(4,5- dimethylthiazol-2-yl)-2,5-diphenyltetrazolium bromide) cell proliferation assay was performed with this cell line as previously described [20, 50, 93]. In brief, HepG2 cells were passaged at 70-80% confluency and the cell suspension was dispensed into a 96 well black, clear bottom falcon plate, aiming to prepare a plate with 20,000 cells/50 µl in each well. Dose response titrations of each compound concentration (200, 100, 75, 50, 20, 10 and 1 µM in 1% DMSO) were performed in triplicate. Media and DMSO (1%) negative controls as well as a positive control (1% v/v Triton X-100, Sigma-Aldrich) were also included in each screen (each of them in triplicate). After addition of compounds, each plate was then incubated for a further 20 h before application of MTT reagent (final 4 h, for a total of 24 h incubation) for assessment of compound cytotoxicity using the MTT assay [63]. Dose response curves were generated in GraphPad Prism 7.02 based on the average absorbance of the three replicates for each concentration point. These data were then used to calculate HepG2 CC_50_ values (the concentration that reduced cell viability by 50% when compared to untreated controls).

### Vitellocyte and egg volume quantification

The total number of *in vitro* laid eggs (IVLEs) produced by compound **33** (3.13 μM) or DMSO treated adult worm pairs were enumerated and fixed in 10% formaldehyde for at least 2 h. Eggs were prepared for laser scanning confocal microscopy (LSCM) visualisation as previously described [20] using DAPI (4’,6-diamidino2-phenylindole, 2μg/ml). Fluorescent microscopic images (10 eggs per treatment) were acquired on a Leica TCS SP8 super resolution laser confocal microscope fitted with a 63X objective (water immersion, 1,75 zoom factor) using the Leica Application Suite X. For each Z-stack, a total of 60 sections were acquired selecting the green (488 nm) and DAPI (405 nm) fluorescent channels for egg autofluorescence and nuclei stain, respectively. Quantification of overall volume (mapped by the green autofluorescence) and content of vitellocytes (DAPI) for individual eggs was performed using IMARIS 7.3 software (Bitplane).

### Quantification of adult worm stem cell proliferation

*In vitro* 5′-ethynyl-2′-deoxyuridine (EdU) labelling of adult worms treated with compound **33** (3.13 µM) or DMSO was performed as previously described [78]. Briefly, adult worms were cultured for 48 h and pulsed with 10 µM EdU for the following 24 h. Following incubation, the worms were fixed, stained and prepared for LSCM. Anterior, posterior and gonadal regions (ovaries for females and testes for males) of both sexes were imaged using a Leica TCS SP5II confocal microscope (40X objective, 1 zoom factor). A Z-stack, containing 60 sections, was generated for each microscopic image for each adult schistosome examined (6 male and 6 female worms for each treatment). Quantitative analyses of DAPI stained non-proliferating nuclei and EdU^+^ nuclei (as representative of all dividing cells) were performed using Imaris software as previously described [15].

### Western blot analysis

Following 72 h post treatment with a sublethal concentration of compound **33** (3.13 µM), adult male worms (n = 20 worms, three biological replicates) were homogenized with a TissueLyser (Qiagen) and total histones extracted using the EpiQuikTM Total Histone Extraction kit (OP-0006, Epigentek) following the manufacturer’s instructions. A total of 2.5-10 µg of each sample was separated by sodium dodecyl sulfate-polyacrylamide gel electrophoresis (SDS-PAGE) through 12% Mini-PROTEAN® TGX™ Precast Gels (4561043, Biorad). After transferring the proteins onto a 0.2 µm Nitrocellulose membrane (Biorad) using a Trans-blot Turbo Midi system (Bio-Rad; Trans-blot turbo protocol - 25V and 2.5A during 3 minutes), the membranes were blocked overnight in 5% fat-free dry milk in Tris-buffered saline (TBS) supplemented with 0.05% Tween 20 (TBST). Following that, the membranes were probed with 1:2000 dilution of anti-H3K4me2 (Abcam, ab32356; Lot GR84714-4) overnight (at 4°C) in 5% fat-free dry milk in TBST. The blot was washed, incubated overnight at 4°C with 1:500 dilution of the secondary antibody (goat anti-rabbit Horseradish Peroxidase coupled antibody - Pierce #31460, Lot HB987318) in 5% fat-free dry milk in TBST. The blot was then developed by incubating with an enhanced chemiluminescence (ECL) substrate (Pierce) followed by CCD camera image acquisition. Acquisition time was adjusted to have maximum exposure without saturation. The detected bands were analysed with ChemiDoc software v4.0.1 with high sensitivity settings.

For sample normalisation, the membranes were stripped by incubation at 50°C for 1 h in 62 mM TRIS-HCl pH 6.8, 2% SDS, 0.8% Beta-Mercaptoethanol, then washed with distilled water several times until complete disappearance of the Beta-Mercaptoethanol smell. The membranes were subsequently probed overnight at 4°C with an anti-H3 Ab (Abcam, ab1791, Lot GR103803-1), 1:1000 diluted in 5% non-fat dried milk in TBST. Detection of H3 signal by secondary antibody, image acquisition and analysis were all performed as described above.

### Statistics

All statistical analyses were performed using a nonparametric Student’s t-test (Mann-Whitney test, two samples) or a two-way ANOVA followed by Least Significant Difference post-hoc correction (more than two samples).

## Acknowledgments

KFH, GP and AB thank the Welsh Government, Life Sciences Research Network Wales scheme and the Wellcome Trust (107475/Z/15/Z) for financially supporting this project. JHK and PAC thank the NIH (GM62437). DL and CG acknowledge the support provided by the framework of the “Laboratoires d’Excellences (LABEX)” TULIP (ANR-10-LABX-41). We thank past and current members of the Hoffmann group and Miss Julie Hirst for contributions to *S. mansoni* lifecycle maintenance. We thank Mr Alan Cookson from IBERS Advanced Microscopy and Bioimaging Suite for assisting with the use of the confocal microscopy facility. We also thank Dr Dylan Phillips and Mr Benjamin J. Hulme for the help during the acquisition and quantification of confocal microscopy images.

## Supporting information

**S1 Figure. Dose response titrations of compounds 15, 16, 33, 35 and 36 against schistosomula**. The five hit compounds were screened against mechanically-transformed schistosomula at 10 µM and lower concentrations (5, 2.50, 1.25 and 0.625 µM). Three independent dose response titrations were performed and each compound concentration was evaluated in duplicate. Each concentration point defines the average mean of the three biological replicates (each of them with two technical replicates). Dose response curves for *S. mansoni* schistosomula phenotype (P) are presented here using GraphPad Prism (mean +/-SE of mean is indicated for each compound concentration). Estimated EC_50_s (and corresponding 95% confidence interval) calculated from these dose response curves are summarised in the table underneath (as well as in **Table 1**). Z′ scores for motility and phenotype for the three screens are reported in **S3 Table**.

**S2 Figure. Compound 33 treated worms causes release of oocytes, spermatozoa and vitelline cells**. After 72 h incubation, representative images of eggs, oocytes (oc), spermatozoa (sp), mature spermatozoa (ms) and vitelline cells (vc) in tissue culture medium of worm pairs treated with DMSO (**A** and **C**) and a sublethal dose of compound **33** (3.13 μM, panels **B, D, E** and **F**) were taken. Images were acquired with Olympus microscope (4x for panels **A** and **B**), 10x for panels **C** and **D** and 20x for panels **E** and **F**)).

**S3 Figure. LSD1 inhibition significantly affects adult female stem cell proliferation**. Female schistosomes treated with 3.13 μM of compound **33** (n = 6) or DMSO controls (n = 6) for 48 h were subsequently labelled with EdU for an additional 24 h. (**A**) - Hand drawing summarising the morphology of *S. mansoni* adult female with representative anterior (yellow box), gonadal (magenta) and posterior region (brown) of untreated (top row, grey) compared to the drug-treated (bottom row, light brown) worms. Scatter plots illustrate the percentage of stem cells present in control (DMSO treated worms, n = 6) versus drug-treated females (3.13 µM of compound **33**, n = 6) in the head region (**B**), the ovary (**C**) and the tail region (**D**). Confocal microscopy, quantification and data visualisation was performed similarly to Fig 5. Standard errors are shown and Mann-Whitney t-test (with **** corresponding to *p* < 0.0001) was subsequently performed to explore a statistical difference between the treatments.

**S4 Figure. Dose response titration of the five hit compounds on HepG2 cells**. Cells were co-cultivated with the five compounds during dose response titrations and subjected to MTT assays. Each titration was performed in triplicate during two independent assays. All absorbance readings are adjusted for background absorbance and the dose-response curve is calculated based on the absorbance reading values from two independent data sets. The absorbance values of the positive (DMSO, 1.25% v/v; x = −2.5) and negative (Triton X-100, 1% v/v final concentration, 0% cell viability value on the y-axis) controls are included. Standard errors of the mean are included on the graph.

**S5 Figure. Computational preparation of the covalent adduct derived from the interaction of compound 33 with the FAD cofactor**. (**A**) - Chemical structure of compound **33**. (**B**) - Stick representation of the chemical structure of the cofactor FAD. (**C**) - Stick representation of the covalent adduct of compound **33** with the flavin ring of the cofactor.

**S6 Figure. Chemical space covered by the library of 39 HsLSD1 inhibitors**. The scatter plot shows the distribution of the calculated logP (cLogP) vs the molecular weight (MW, g/mol) of the 39 compounds under study in this investigation. Each compound is shown as orange dot where the five most active compounds are shown in green. The reference compound (compound **1**) of this family of LSD1 inhibitors and the two more closely related derivatives (compounds **2** and **3**) are labelled for comparison to the five most active anti-schistosomal compounds presented in this study.

**S1 Table**. Chemical structures of the 39 compounds included in this study.

**S2 Table**. Z′ values for both phenotype and motility of the Roboworm screens performed on the 39 compounds.

**S3 Table**. Z′ values for both phenotype and motility of the Roboworm screens performed on the titration of the five selected compounds (compounds **15, 16, 33, 35** and **36**).

**S4 Table**. List of the structure, the docking score and EC_50_ values (on schistosomula and adult worms) of the five most active compounds.

**S1 Movie**. Video of *S. mansoni* drug treated (compound **33**, 3.13 µM, right-hand side) and control (DMSO, left-hand side) worm pairs after 72 h incubation in tissue culture wells. Notice the lack of parasite attachment and the presence of cellular material within the compound treated well.

**S2 Movie**. Serial optical sections of DAPI-stained, *S. mansoni* egg. Comparison between the drug treated (compound **33**, 3.13 µM, right-hand side) and the negative control (DMSO, left-hand side) is provided.

**S3 Movie**. Serial optical sections of *S. mansoni* adult male worm stained with DAPI and EDU. In these series of optical sections, three different anatomical regions (anterior region, gonadal system and posterior region, from left to right) of the worm were observed. Comparison between the negative control (DMSO, first row) and the drug treated (compound **33**, 3.13 µM, second row) parasites is provided.

**S4 Movie**. Serial optical sections of *S. mansoni* adult female worm stained with DAPI and EDU. In these series of optical sections, three different anatomical regions (anterior region, gonadal system and posterior region, from left to right) of the worm were observed. Comparison between the negative control (DMSO, first row) and the drug treated (compound **33**, 3.13 µM, second row) parasites is provided.

